# Knowledge, attitudes, and practices of para-veterinary workers about ticks and tick-borne diseases in three provinces of Pakistan

**DOI:** 10.1101/2025.08.05.668794

**Authors:** Abrar Hussain, Sabir Hussain, Mahvish Rajput, Muhammad Sohail Sajid, Olivier Sparagano, Nohra Mateus-Pinilla, Rebecca L. Smith

## Abstract

There is a high prevalence of tick infestation in Pakistani livestock, affecting more than 45% of the population of more than 200 million small and large ruminants. Most livestock farmers seek assistance from para-veterinary workers, who fall under the definition of Veterinary Paraprofessionals (VPPs), according to the World Organization for Animal Health (WOAH). There is a shortage of information concerning the awareness of these para-veterinary workers regarding tick control and management. This study aims to bridge this critical knowledge gap by conducting a cross-sectional survey that evaluates the knowledge, attitudes, and practices of para-veterinary workers about tick-borne diseases (TBDs) in Pakistan. Between March and August 2023, we conducted a web-based survey among para-veterinary workers recruited via email, text message, and face-to-face conversations. Poisson regression was used to identify factors associated with knowledge, attitude, and practice (KAP) scores related to TBDs. We received 118 responses from three provinces; only 27.9% (*n* = 33) responded that they had attended workshops related to ticks and TBDs. Attending workshops was associated with higher KAP scores. All section scores were correlated, and higher knowledge scores were significantly associated with lower odds of tick exposure. Our findings suggest that workshop attendance is important in increasing overall awareness and promoting better practices regarding TBDs.

## 1. Introduction

Pakistan has a huge livestock sector, contributing over 60% of the value of the agriculture sector and almost 11% of the country’s GDP (Rajput et al., 2023). The livestock sector mainly includes small farm holders; about 30-35 million people rely on it for their livelihood, producing 80% of Pakistan’s total milk (Khan et al., 2022). According to a livestock survey of Pakistan, there are more than 200 million ruminants, including 48 million cattle and 40 million buffalo, owned by rural families (Hussain et al., 2022a). Among this population of ruminants, the prevalence of tick infestation has been reported to be 34.83% (buffaloes), 57.11% (cattle), 51.97% (sheep), and 46.94% (goats) (Khan et al., 2022). The issue of tick control in Pakistan has gained importance over the past few decades due to the introduction of exotic cattle breeds for the subsistence or commercial dairy sectors. These imported breeds are more susceptible to ticks and tick-borne diseases (TBDs) than indigenous breeds, such as Sahiwal cattle, which deter tick attachment by twitching their skin (Jabbar et al., 2015). Ticks pose a significant threat to human and animal health through infestation and the transmission of viral, bacterial, and protozoal pathogens, collectively referred to as TBDs (Ogden et al., 2021). The global impact of ticks and TBDs is steadily increasing, escalating burdens on human and animal health (Boulanger et al., 2019).

*Hyalomma* and *Rhipicephalus* tick species represent significant hazards to livestock production in Pakistan, predominantly contributing to the incidence of babesiosis, theileriosis, and anaplasmosis in ruminants (Karim et al., 2017). *Rhipicephalus microplus*, commonly known as the cattle tick, is a proficient vector for *Babesia bovis, B. bigemina*, and *Anaplasma marginale*, causative agents of tick fever affecting livestock in Pakistan and worldwide. *Hyalomma* species, on the other hand, act as known vectors for *Theileria annulata*, an animal disease resembling malaria (Jabbar et al., 2015)*. Hyalomma* tick bites can also transmit Crimean-Congo Hemorrhagic Fever (CCHF), a zoonotic viral TBD (Mourya et al., 2019). Zoonotic TBDs, especially anaplasmosis, babesiosis, and ehrlichiosis, can be transmitted to livestock workers through the crushing of carrier ticks by bare hands or through contact with infected blood or tissues of animals during or after slaughter (Ahmed et al., 2021b). Individuals at higher risk of contracting CCHF in endemic regions include farmers, para-veterinary workers, slaughterhouse workers, veterinarians, and healthcare professionals (HCPs) (Ahmed et al., 2021a). The saliva of both these ticks can also lead to skin lesions and systemic reactions in humans.

Smallholder farmers, predominantly depend on para-veterinary workers for assistance concerning the health of their animals (Hussain et al., 2021b). This reliance places para-veterinary workers in a crucial position in managing and controlling TBDs. It is imperative to evaluate these para-veterinary workers’ knowledge, attitudes, and practices to gauge their awareness of tick-borne disease control and address deficiencies in their knowledge through training. To accomplish this objective, we conducted a cross-sectional study in various regions of Pakistan to assess para-veterinary workers’ knowledge, attitudes, and practices regarding ticks and TBDs.

## 2. Material and Methods

### 2.1. Survey design and participant recruitment

A web-based survey focusing on knowledge, attitudes, and practices related to ticks and TBDs relevant to Pakistani livestock was developed within RedCap (Research Electronic Data Capture), a tool hosted at the University of Illinois, Urbana-Champaign (Harvey, 2018). RedCap offered several essential features, including a user-friendly interface for validated data entry, audit trails to track data manipulation and export activities, and data download procedures with automated de-identification capabilities. The survey questions were pre-tested by researchers familiar with the study topic. Based on their suggestions, complex terminologies were removed, and additional questions were incorporated. The target study population consisted of all individuals providing animal health services in Pakistan, excluding veterinarians. These para-veterinary workers include veterinary assistants (VAs) with a two-year Livestock Assistant Diploma and artificial insemination (AI) technicians with a six-month diploma. Both fall under the definition of Veterinary Paraprofessionals (VPPs), as defined by the World Organization for Animal Health (WOAH) (“Veterinary Workforce Development - WOAH,” 2022), and practitioners without formal veterinary education who sometimes use herbal medicine, come under the definition of Community animal health worker (CAHW) (“Veterinary Workforce Development - WOAH,” 2022). This paper will use para-veterinary workers for both VPP and CAHW. The questionnaire encompassed four distinct categories: 1) Demographics (age; sex; designation as being VA, AI technician, or no-certification; experience in years; and any past training received related to TBDs); 2) Knowledge (understanding of ticks and TBDs in animals and zoonotic implications of TBDs); 3) Attitudes (level of concern regarding ticks and TBDs, interest in attending workshops focused on TBDs); and 4) Practices (use of diagnostic tests and treatment strategies for common TBDs, efforts to educate farmers about tick and TBD control in animals and zoonotic risks associated with TBDs, and personal protective measures used during and after farm visits). The questions in the survey were primarily presented in a multiple-choice format, with some short-answer questions. Factual responses were categorized as “correct” or “incorrect” based on information widely available through reputable sources such as the Centers for Disease Control and Prevention (CDC) website (“Diseases Transmitted by Ticks | Ticks | CDC,” 2019.), and previously published articles (Hussain et al., 2022b; Hussain et al., 2021a). The questionnaire was available in English and Urdu, the national language of Pakistan and the 11th most widely spoken language globally, and is understood to some extent by almost the entire population of Pakistan (“Information for providers of families/parents who speak Urdu,” 1996). Almost all the para-veterinary workers have at least 10 years of education in Urdu and 3 years in English **(S1)**.

A convenience sampling strategy was employed to target para-veterinary workers in Pakistan for this survey. The online survey was disseminated through various channels, including social networking platforms (sending links to the online survey through WhatsApp and Facebook) and professional networks (asking veterinarians and participating para-veterinary workers to recruit additional participants by word of mouth). The study’s recruitment strategy, which included disseminating the survey across various social media platforms, aimed to mitigate network bias by reaching a broader demographic beyond the immediate professional and social circles connected to the initial respondents and researchers. For online responses, the questionnaire remained accessible from April 2023 to August 2023, containing the consent form on the first page, allowing participants to submit their responses within this time frame. Professional network recruitment included face-to-face meetings with one of the researchers from May 2023 to June 2023.

### 2.2. Ethical statement

The study received approval from the National Bioethics Committee of Pakistan, and the University of Illinois Urbana-Champaign Institutional Review Board approved the study protocol. (protocol # 23575).

### 2.3. Theoretical Framework

The model was developed based on the theory of knowledge, attitudes, and practices (KAP) (“Advocacy, communication and social mobilization for TB control: a guide to developing knowledge, attitude, and practice surveys,” 2008). Knowledge was assessed using 11 questions, attitudes with eight questions, and practices with nine questions, with each domain further categorized into four aspects **(Figure 1).** A scoring system was implemented by assigning points to correct responses, which were then summed up to generate a total score for each section per respondent **(S2)**.

**Fig. 1.**
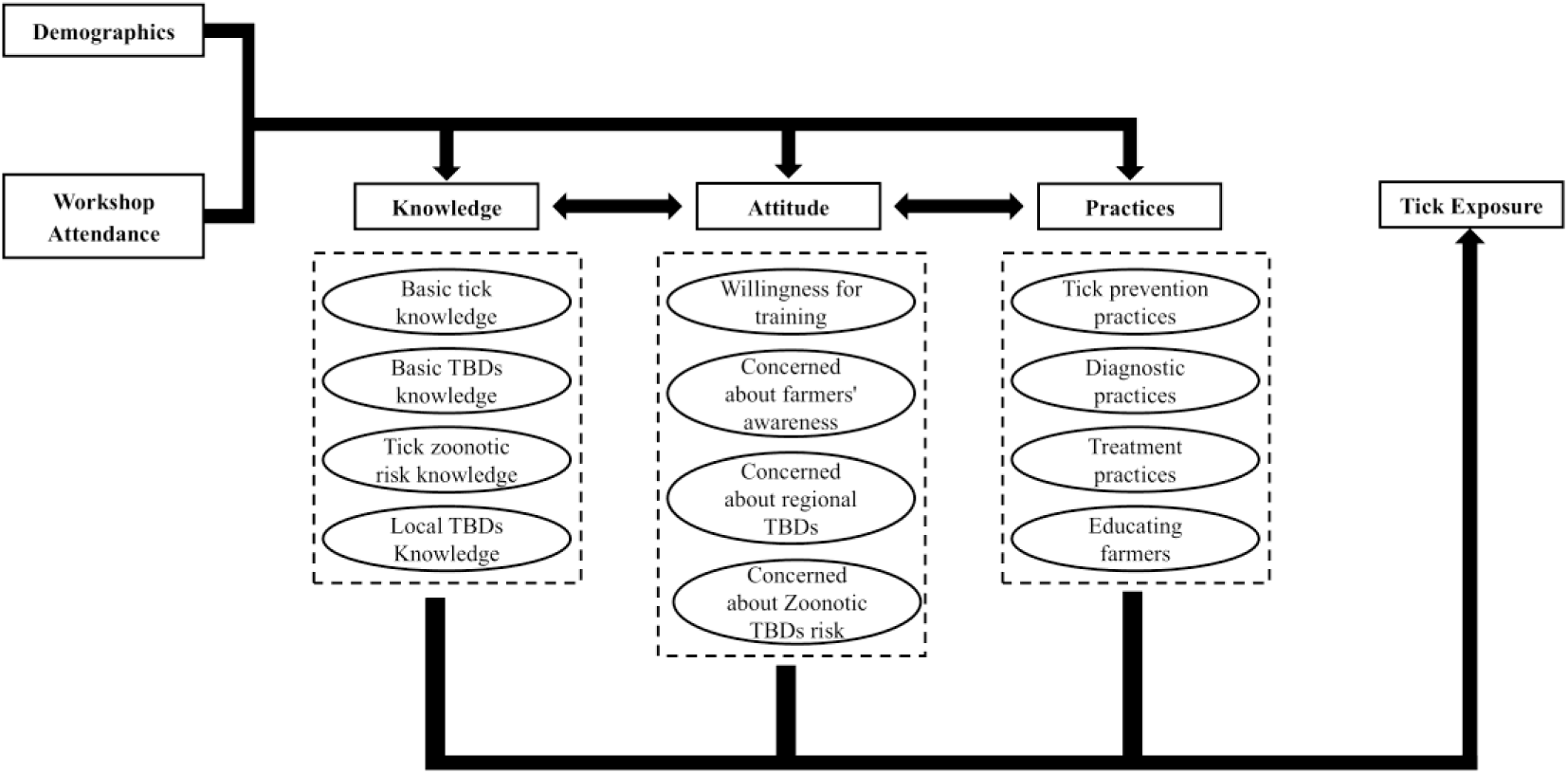
The theoretical framework illustrates the relationships among knowledge, attitudes, and practices (KAP) regarding ticks and tick-borne diseases, including potential influencing factors and expected outcomes.

The model hypothesizes that workshop participation influences knowledge, attitudes, and practices. Additionally, demographic factors may also impact behaviors, and higher scores in knowledge, attitudes, and practices are expected to reduce personal exposure to ticks (Figure 1). **(Figure 1).**

### 2.4. Statistical analysis

Data were analyzed in R Studio version 4.1.3 (“R: The R Project for Statistical Computing,” 2019.). The outcome variable was the total score for each section, calculated based on the number of correct responses. Poisson regression was employed to identify significant predictors of higher scores. Logistic regression was used to examine the relationship between scores and tick exposure for each section separately, while Spearman’s correlation was applied to assess the association among scores across different sections.

## 3. Results

During the six-month survey period, a total of 159 responses were obtained. After excluding 41 incomplete responses, the final analysis was conducted with 118 complete responses.

### 3.1. Demographics

The survey encompassed 22 districts from three provinces in Pakistan **(Figure 2)**; these provinces contain approximately 75% of the country’s total human population. All respondents were male, 72% being veterinary assistants, 21%AI technicians, and 7% without a formal certification. Most respondents (62.1%) reported doing field practice (as opposed to clinical, 5%, or mixed practice, 29%). Among the participants, the majority fell within the age range of 35 to 54 years (58%) and had professional experience of one to ten years (51.7%). All respondents reported dealing with tick-related health issues in animals, encountering a monthly average of 17 tick-borne disease cases (range: 2, 60). Additionally, 27.9% (n = 33) of the 118 participants indicated attending workshops and receiving training focused on ticks and TBDs **(Table 1)**.

**Fig. 2.**
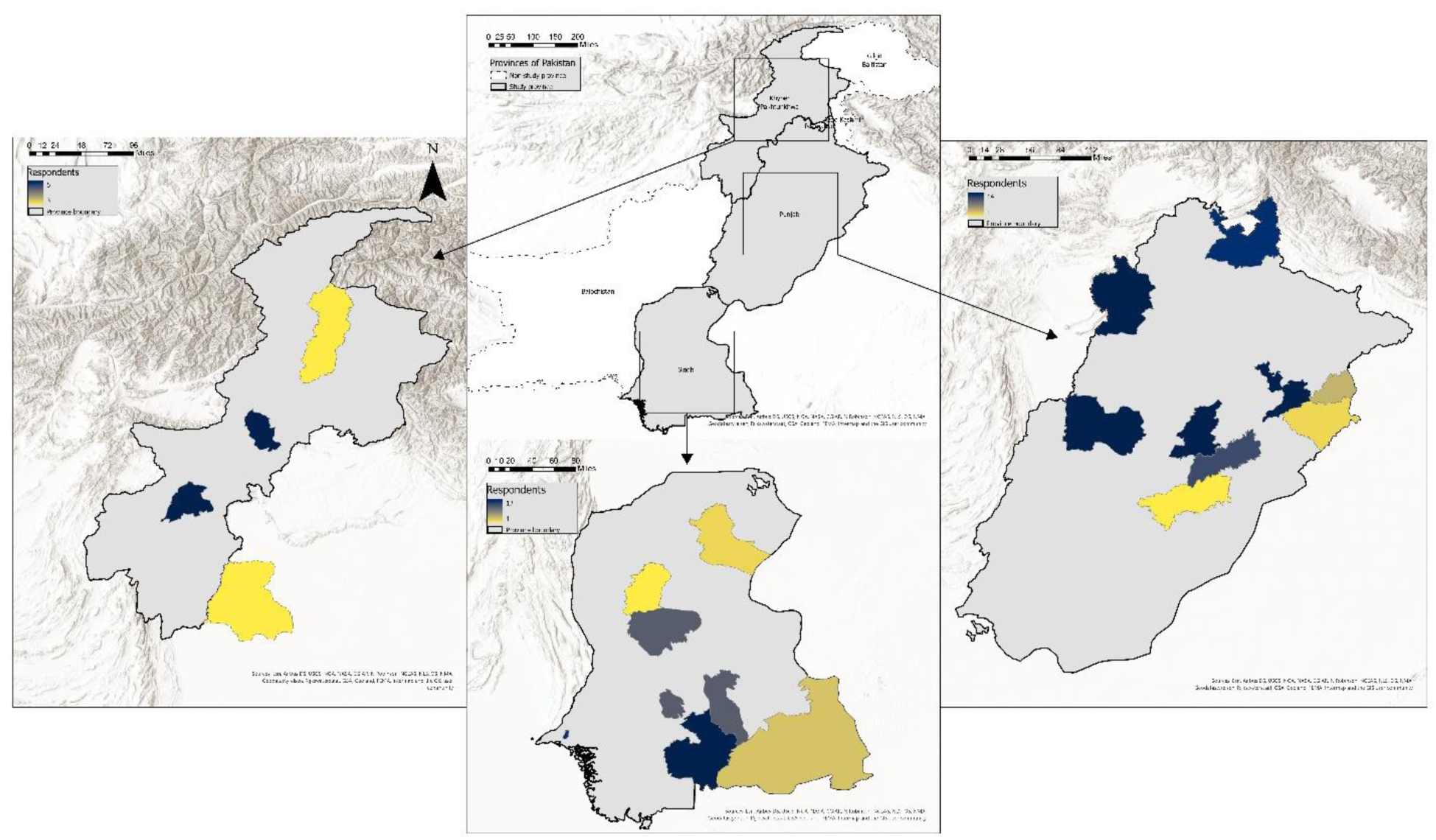
The geographical distribution of study locations in Pakistan displays the number of para-veterinary worker respondents from different districts.

**Table 1.**
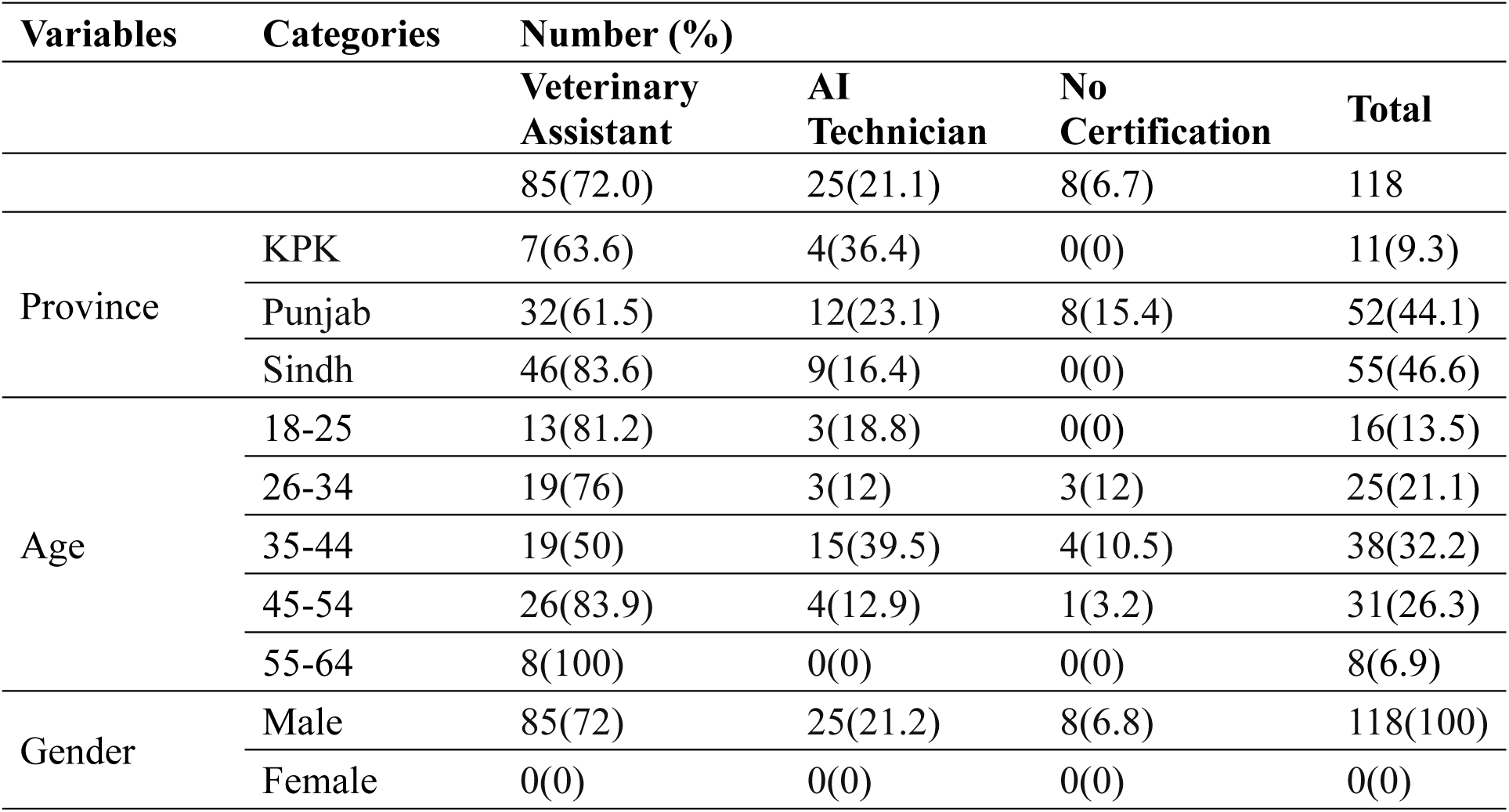
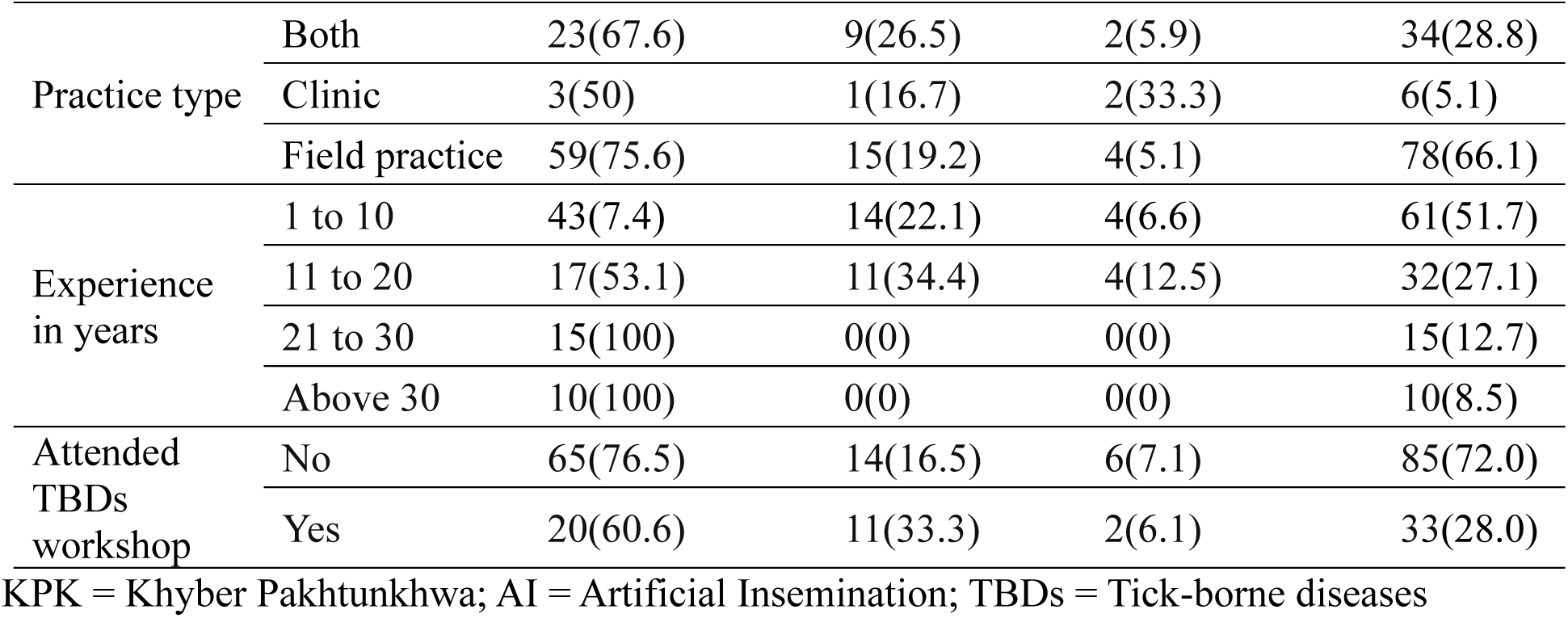
Demographics of respondents from differences by designation.

### 3.2. Knowledge

The most frequently reported sites of tick bites were the udder, inner thighs, and ears of animals. A significant majority (80.5%) identified cattle and buffalo as the animals most infested with ticks, with 60.1% indicating that ticks prefer habitats such as cracks and crevices. Despite this, 79.6% of respondents were unfamiliar with the names and identification of common tick species in their region, and 52.5% believed that the sensation of a tick bite was felt immediately.

While 77.1% acknowledged that ticks could transmit zoonotic diseases, 61.8% could not specify which zoonotic TBDs are prevalent; 41.5% identified babesiosis, and 36.4% identified theileriosis as endemic in their areas, despite both diseases being endemic in the entire study area. However, nearly 75% expressed high concern over TBDs in their localities (**Figure 3**).

**Fig. 3.**
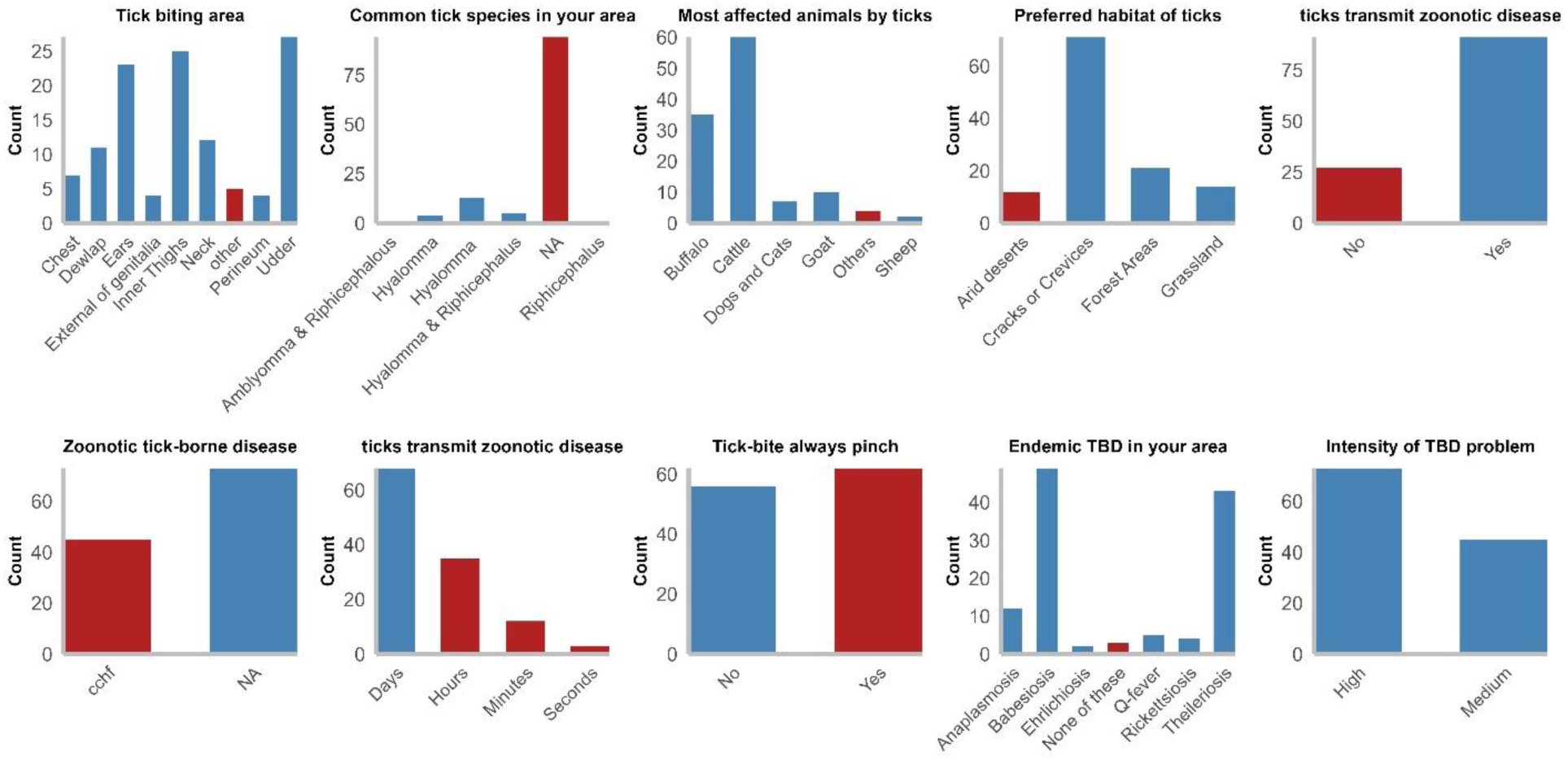
Distribution of responses to knowledge-related survey questions assessing para-veterinary workers’ awareness and understanding of ticks and tick-borne diseases (blue color = correct, red color = incorrect).

### 3.3. Attitudes

When asked about the necessity of training on TBDs, 92.4% of respondents expressed support, with 66.1% preferring in-person training. Nearly 97% of respondents believed raising awareness about TBDs among farmers is critically essential. They identified a lack of farmer concern and the cost of acaricides as major obstacles to controlling TBDs. According to 95% of respondents, animals in their area are at significant risk of TBDs, and 90% reported that farmers are consistently at risk of tick bites (**Figure 4**).

**Fig. 4.**
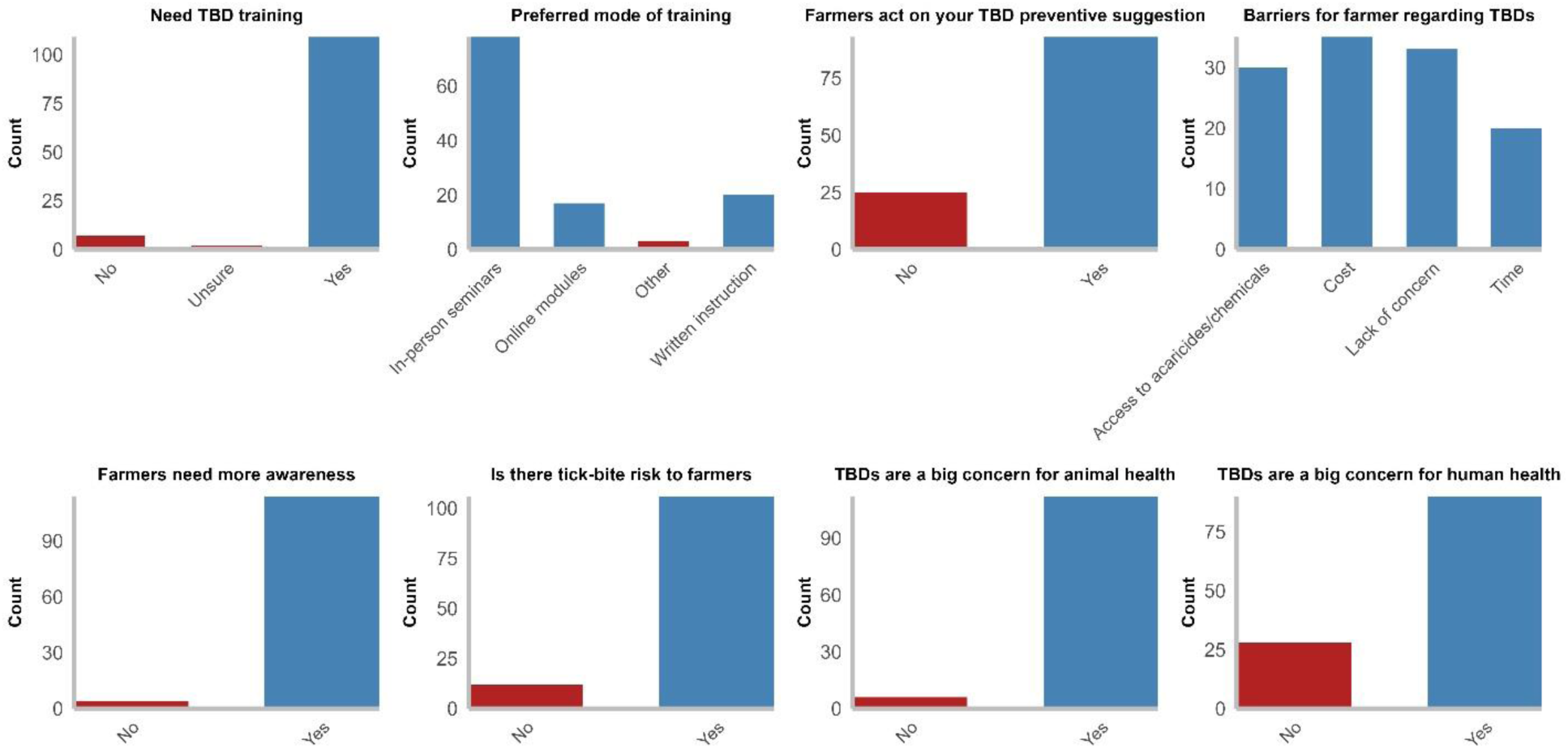
Distribution of responses to attitudes-related survey questions evaluating para-veterinary workers’ perceptions, concerns, and beliefs regarding ticks and tick-borne diseases (blue color = correct, red color = incorrect).

### 3.4. Practices

Approximately 38% of para-veterinary workers reported using diagnostic tests for TBDs; however, none could specify any diagnostic test by name. A majority, 80.1%, were optimistic about their role in educating farmers about TBDs, and 40.1% reported recommending acaricides as an effective method for tick control. About 46.6% indicated that farmers had reported tick bites at least once during their careers, and 37.1% of respondents had personally experienced a tick bite at least once. However, 66.1% did not employ protective measures while handling tick-infested animals and failed to check themselves for ticks after visiting livestock farms (**Figure 5**).

**Fig. 5.**
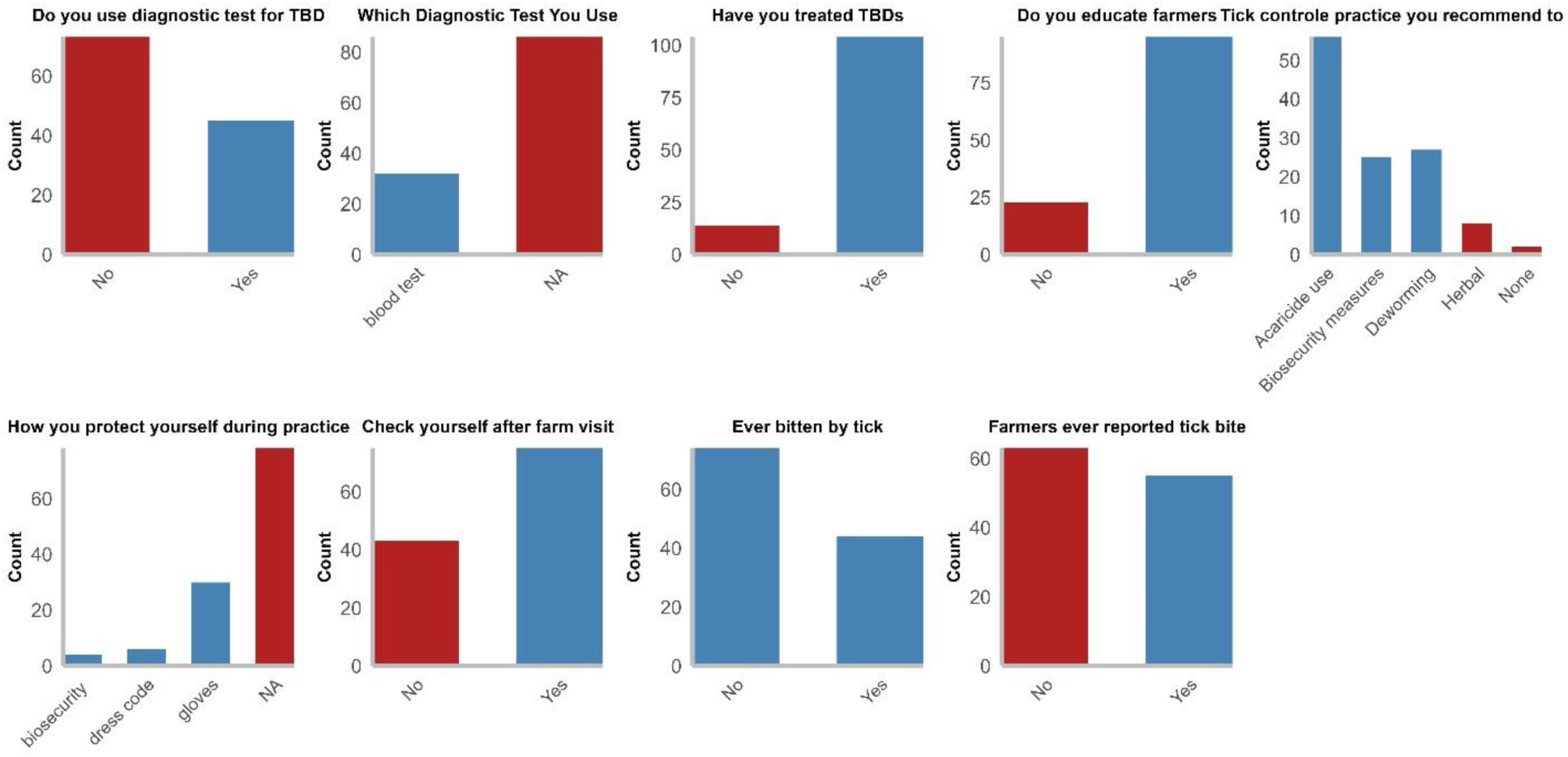
Distribution of responses to practice-related survey questions assessing para-veterinary workers’ reported behaviors and preventive measures against tick-borne diseases (blue color = correct, red color = incorrect).

### 3.5. Predictors of knowledge, attitudes, and practices scores

Practice area (province), age, designation, practice type, professional experience, and workshop attendance were included as potential covariates in the Poisson regression model. No variables were significantly related to any of the scores among demographics. Workshop attendance was the most influential factor contributing to a higher knowledge score (OR = 1.23, 95% CI 1.13 – 1.34), attitudes score (OR = 1.15, 95% CI 1.05 – 1.26), and practice score (OR = 1.41 95% CI 1.18 – 1.70) **(Table 2)** (**Figure 6**).

**Fig. 6.**
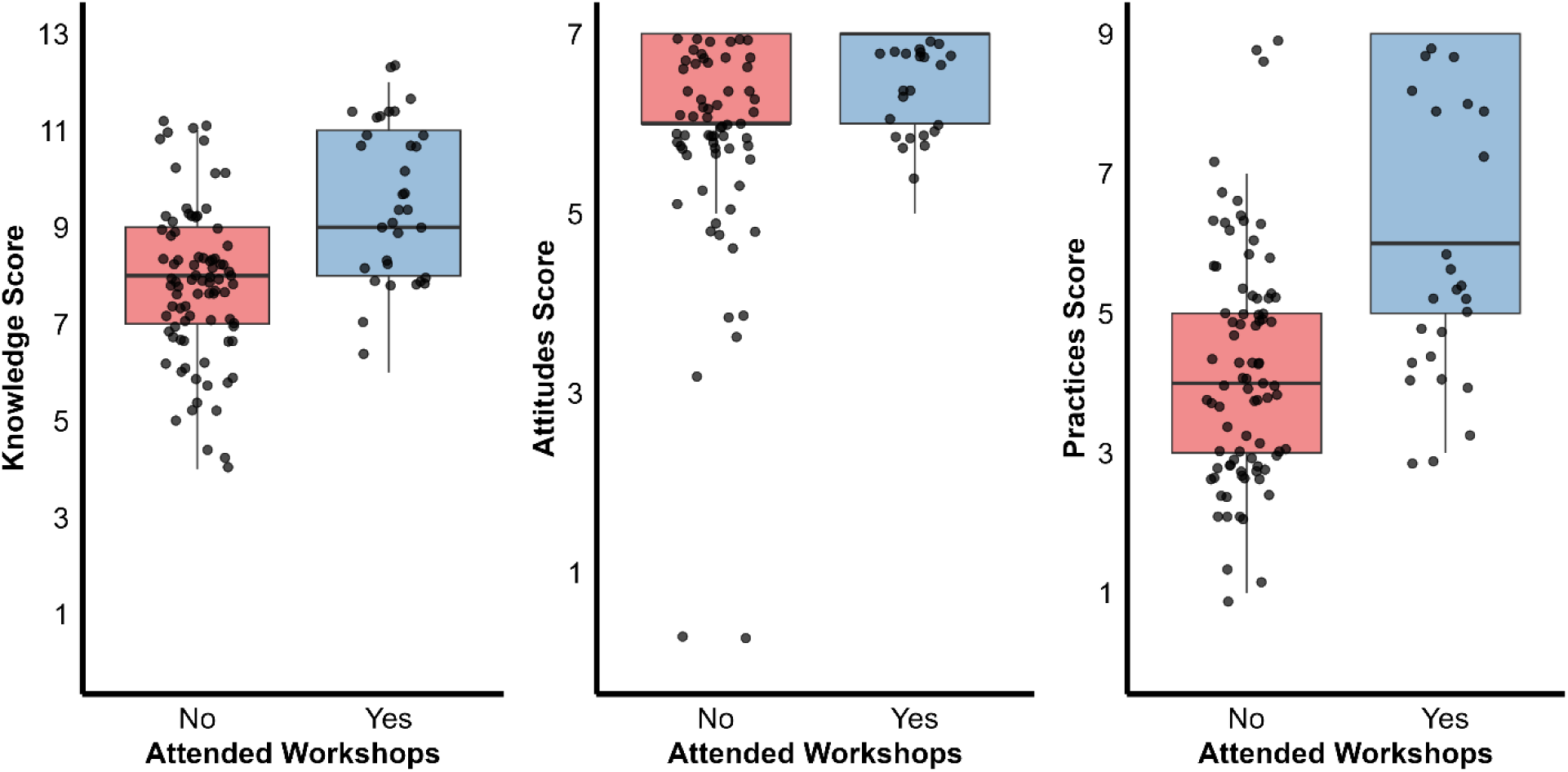
Comparison of total scores for knowledge, attitudes, and practices (KAP) on ticks and tick-borne diseases among para-veterinary workers, stratified by workshop attendance status.

**Table 2.**
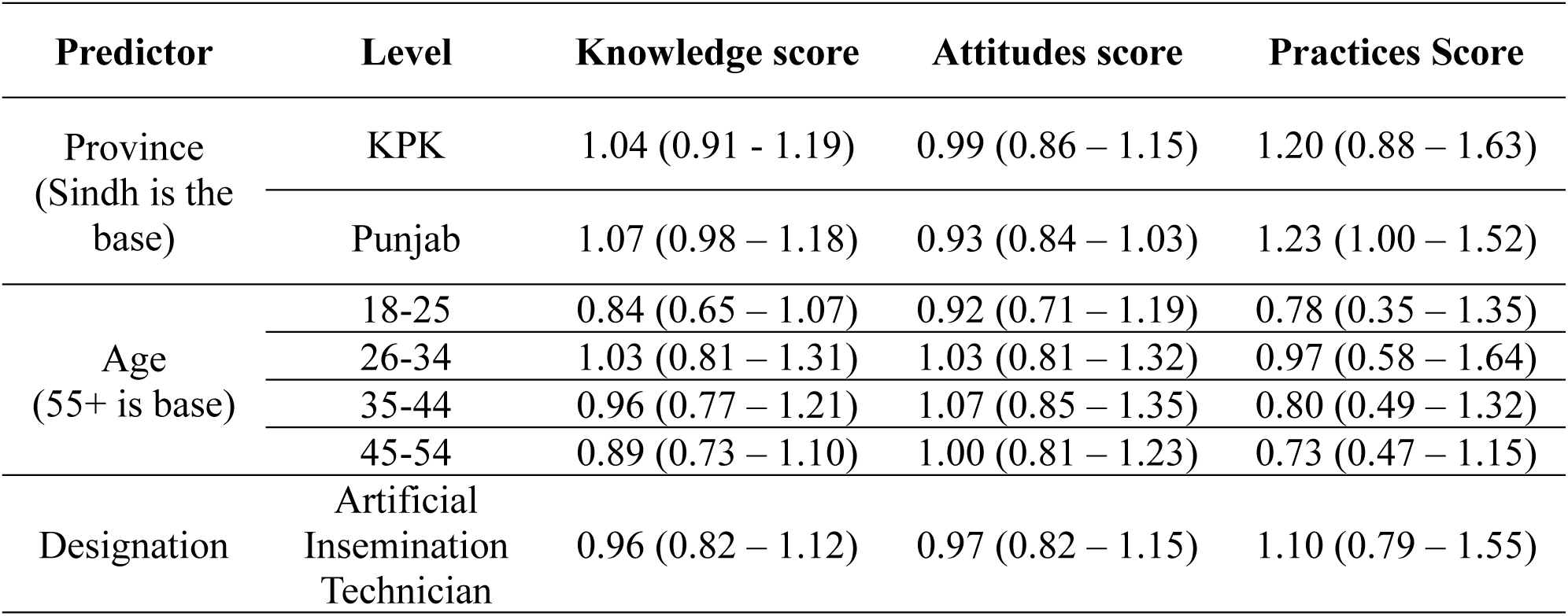
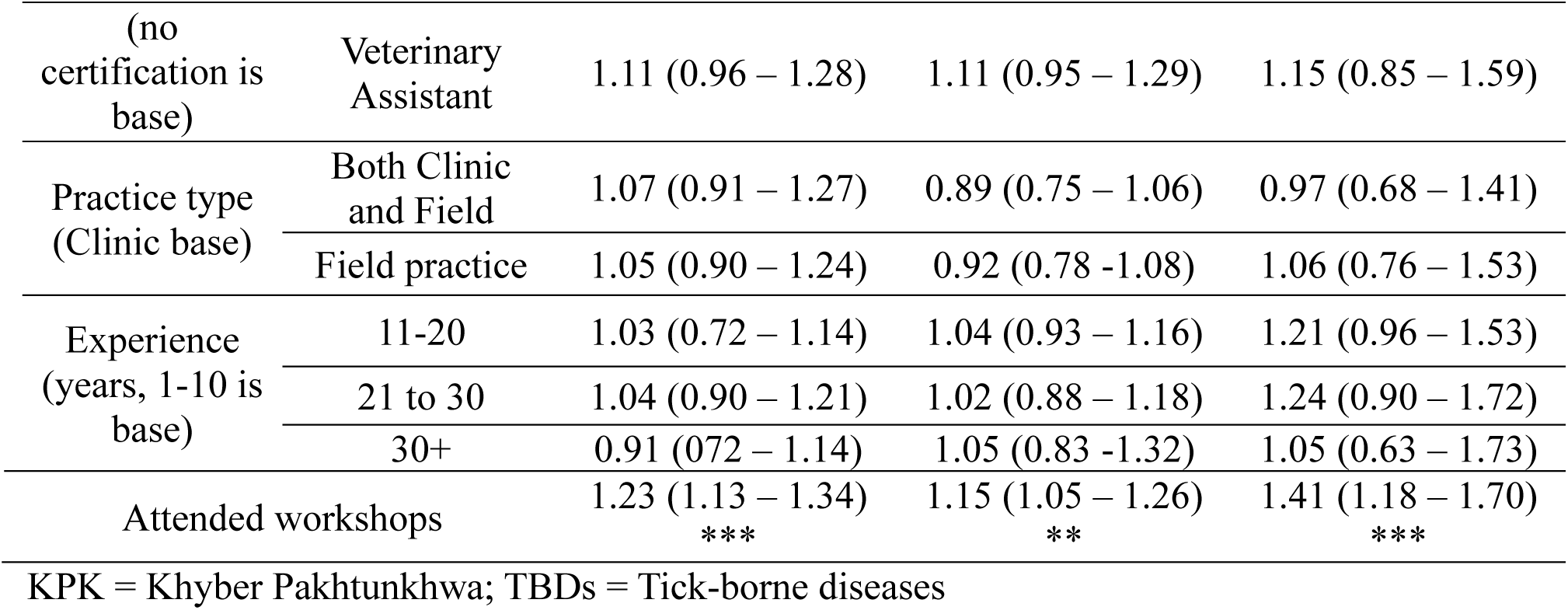
Poisson regression model of predictors associated with tick knowledge, attitudes, and practices score.

### 3.6. Predictors of tick exposure

All three section scores were correlated with each other **(Figure 7)** and were used as potential covariates in three logistic regression models. While higher knowledge, attitudes, and practice scores were generally associated with lower odds of tick bites, only the knowledge score demonstrated a statistically significant effect **(Figure 8)**. Specifically, for each one-unit increase in the knowledge score, the odds of experiencing a tick bite decreased by approximately 39%.

**Fig. 7.**
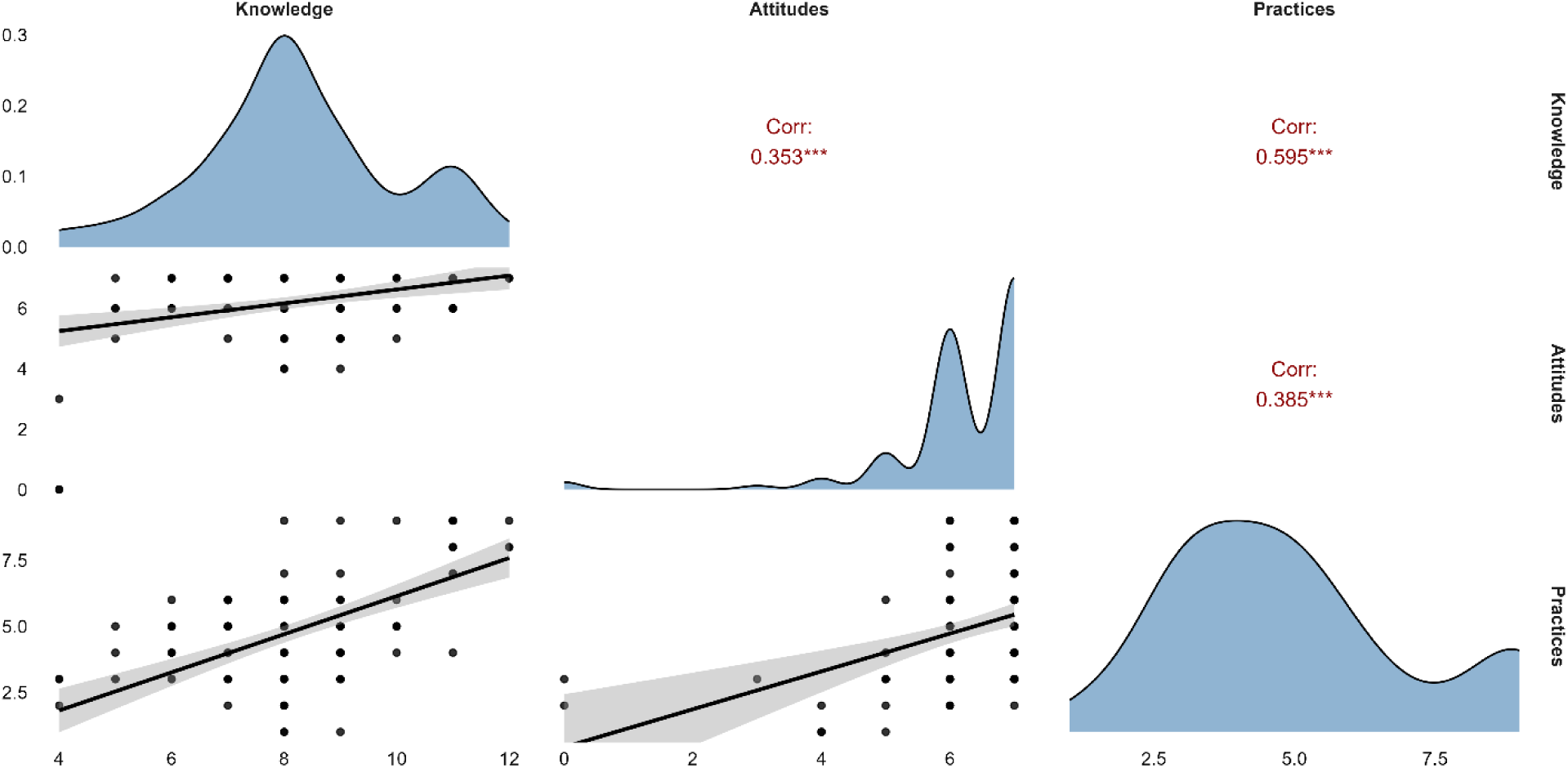
The Correlation matrix illustrates the relationship between knowledge, attitudes, and practice scores among para-veterinary workers, highlighting potential interdependence.

**Fig. 8.**
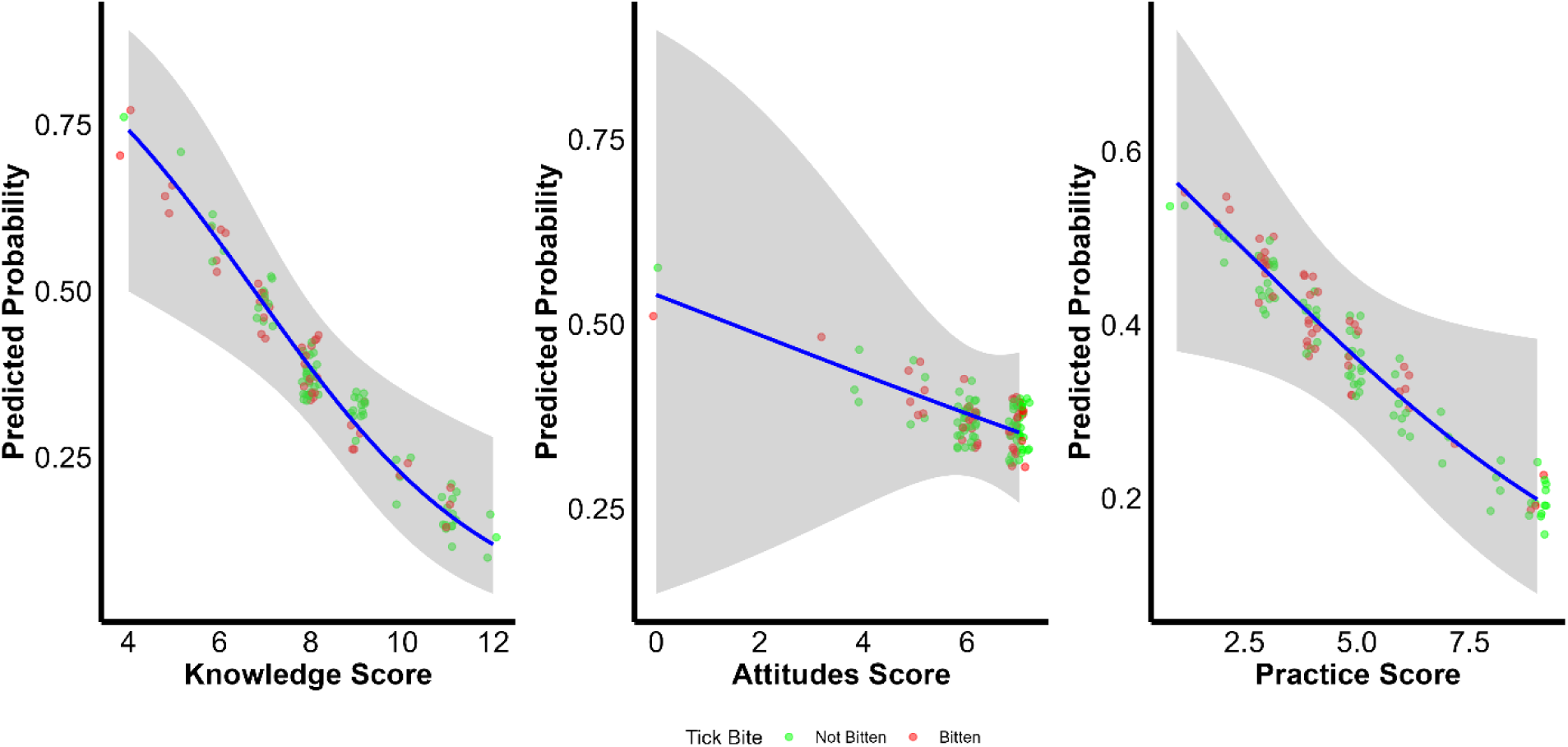
Logistic regression results show the association of knowledge, attitudes, and practice scores with the likelihood of experiencing a tick bite among para-veterinary workers.

## 4. Discussion

While we found that para-veterinary workers in Pakistan were concerned about tick-borne diseases and aware of the high levels of tick infestation in the livestock they treated, there were gaps in their knowledge, attitudes, and practices, particularly among those who did not update their knowledge through workshops. Given the central role of para-veterinary workers in providing veterinary care and animal health information in Pakistan, understanding these gaps and how they might be addressed is crucial.

Most responses were from Punjab and Sindh, Pakistan’s two most populous provinces, which also have a significant proportion of the country’s ruminant population. Workshop attendance was notably higher in Punjab (50.0%) than in the provinces of Sindh (7.2%) and KPK (27.5%). This could be attributed to the comparatively elevated literacy rate in Punjab, along with its larger population of animals (Khan et al., 2022). These workshops are offered freely by the livestock departments of the respective provinces through gatherings at different civil veterinary hospitals. There was no significant difference between veterinary assistants (VAs), AI technicians, and those with no certification for attending TBD workshops.

The results of the Poisson regression model depicted that TBD workshop attendance was associated with higher knowledge, attitudes, and practice scores, suggesting that the workshops may have a constructive role in enhancing tick-related practices among para-veterinary workers. While workshop attendance improved awareness, knowledge gaps regarding tick identification remained prevalent. Approximately 80% of respondents, including 28% who had attended TBD workshops, lacked familiarity with common tick species in Pakistan. This gap may arise from recognizing ticks as general pests without understanding species-specific disease transmission. Additionally, there appears to be a lack of awareness regarding these tick species’ common and scientific names. This underscores a need for comprehensive education about ticks and suggests room for improvement in the content and quality of the workshops according to the target audience. The Punjab Livestock Department has recently initiated disease surveillance and anti-tick campaigns, encompassing mobile veterinary disease diagnostic lab visits and capacity-building programs for veterinarians and para-veterinary workers (“Services | Livestock Punjab,” 2018.). Such education would greatly benefit smallholder farmers, who rely on para-veterinary workers for tick education. A study conducted in Pakistan suggested that it is possible to expand extension programs by using existing government resources, but the main challenge lies in making sure field personnel, such as para-veterinary workers, develop the skills to reach out to farming communities (Warriach et al., 2018). Prior research focusing on farmer-targeted knowledge transfer initiatives in Africa demonstrated tangible enhancements in animal health (Bell et al., 2005; Grace et al., 2008; Stringer et al., 2011).

Effective knowledge exchange programs must cater to the target audience, align with the local context, and ensure accessibility, acceptability, and comprehensibility to ensure knowledge retention. Our study indicated that 66.1% of para-veterinary workers preferred in-person meetings to learn about ticks and TBDs. A study conducted in India yielded similar findings, indicating that the dissemination of posters and leaflets did not notably enhance farmers’ knowledge or partially trained workers, with in-person meetings proving more effective (Hopker et al., 2018). Including visual aids can enhance an educational program; however, the concept of “visual literacy,” which pertains to the reader’s ability to interpret visual materials, must be considered. It is advisable to pilot such materials to ensure their efficacy in conveying the intended information (Domínguez Romero and Bobkina, 2021; Eutsler, 2021). Considering the ongoing anti-tick campaign by the Punjab Livestock Department, a notable emphasis on visual literacy is warranted, particularly for para-veterinary workers who need to identify the most prevalent disease-transmitting tick species. Consequently, printed materials may be best paired with in-person training sessions to educate para-veterinary workers about ticks and TBDs, with the same training serving as a refresher for diagnosing and preventing tick-borne infections.

Among our respondents, one-third reported experiencing at least one tick bite during their careers. A higher knowledge score was significantly associated with lower odds of tick exposure, indicating that increased awareness may contribute to better preventive practices against TBDs. This finding aligns with a recent study, which demonstrated that individuals with extensive knowledge of ticks had a lower risk of contracting TBDs, even in regions with high tick prevalence. (Fang et al., 2024). Volunteer bias may still be present since the respondents who chose to participate are likely more engaged or have stronger opinions about the subject matter. More than half of the respondents never used gloves or any protective measure while handling tick-infested animals or visiting tick-infested livestock farms; while gloves themselves may not prevent tick bites, we used the question about gloves as a proxy for Personal protective equipment (PPE) overall. Para-veterinary workers are at a heightened risk of tick exposure due to their direct interactions with animals and visits to tick-prone environments for on-site care provision (Kinnunen et al., 2022). Our findings of their lack of protective practices highlight the evident necessity for more comprehensive education regarding tick-borne zoonoses and preventive measures, especially given the lack of available vaccines for different tick-borne infections, including infections caused by *Coxiella burnetii, Francisella tularensis*, and rickettsioses (Wölfel et al., 2017).

Para-veterinary workers are crucial in bridging the gap between animal and human health by educating farmers about ticks and TBDs (Wohl and Nusbaum, 2007). Our findings indicate that over 80% of para-veterinary workers reported engaging in raising awareness among farmers about the impact of TBDs. However, many do not update their knowledge and practices through workshops, which may limit the effectiveness of the awareness they create in preventing TBDs. Additionally, nearly half of the respondents reported knowing farmers bitten by ticks, highlighting a significant risk among this group. Therefore, it is essential to emphasize the need for ongoing education among para-veterinary workers regarding the role of ticks as vectors of zoonotic diseases. Strengthening their knowledge and capacity to educate farmers could help reduce the occupational risk of zoonotic TBDs while enhancing awareness and preventive measures at the livestock farming level.

A recent study indicated that para-veterinary workers are the primary source of tick-related information for farmers and animal owners (Hussain et al., 2021). This differs from other countries, where veterinarians predominantly fulfill this role (Evason et al., 2021). However, given the relatively limited number of veterinarians in Pakistan compared to the substantial population of ruminants (approximately 200 million), para-veterinary workers assume a critical role in addressing animal healthcare needs. India faces a similar situation, where para-veterinary workers are primarily responsible for first aid and general dispensing (Kumar and Meena, 2021). However, we found that these para-veterinary workers often lack the skills to inform farmers of TBD’s current risk to their animals and the best ways to protect themselves. Thus, improving the training of para-veterinary workers becomes imperative to enhance farmer awareness about ticks and tick-borne zoonoses, ultimately leading to more effective preventive measures against ticks.

Within our respondent cohort, 6.8% self-reported as unregistered animal health practitioners who lack formal training. During the recruitment process, many unregistered practitioners declined participation in our study (data not shown due to lack of consent), suggesting that the actual count of unregistered animal health practitioners is likely higher than our participant data portrays. These para-veterinary workers without certification may lack access to or awareness of the training available to the other groups; although there was no significant difference in scores or workshop participation for this group, the small sample size limited our power to detect such a difference. There may be a need for attention from bodies such as the Pakistan Veterinary Medical Council, the Livestock Department, and other governmental authorities to ensure that relevant trainings are offered to all the working groups in the animal health community and that the unregistered practitioners feel welcome in those training.

This study had some limitations, mainly due to the convenience sampling process of the participants. However, our sample did show a relatively good representation of different geographical regions and experience levels. In conclusion, this study bridges a critical knowledge gap by evaluating para-veterinary workers’ knowledge, attitudes, and practices concerning TBDs and tick-related risks in various regions of Pakistan. The survey identified the positive impact of attending tick-related workshops on overall awareness and practices. However, gaps in knowledge about tick species and zoonoses persist, emphasizing the need for continuous training tailored to this audience. Enhancing their education could help para-veterinary workers mitigate zoonotic risks for themselves while improving livestock health and welfare.

## Data Availability

The datasets analyzed during the current study are available in the University of Illinois Urbana Champaign data repository. Here is the reviewer’s link: https://databank.illinois.edu/datasets/IDB-0452634?code=YetT02lG_q5LvZhHAuhTZGZHbBclDRB769F-MkdICFE

Here is the DOI that will be active after publication: https://doi.org/10.13012/B2IDB-0452634_V1

## Supporting information

Codebook from REDCap

Scoring criteria

## Acknowledgments

We thank all the para-veterinary workers for their participation. We thank the veterinarians, especially Dr. Asif Maqsood and Dr. Fazal-e-raziq Yousufzai, who helped us connect with para-veterinary workers and data collection.

## List of Supplementary files

**S1 File.** Codebook from REDCap

**S2 File**. Scoring criteria

