## Supplementary material for "Knowledge, attitudes, and practices of para-veterinary workers about ticks and tick-borne diseases in three provinces of Pakistan": Codebook from REDCap

### Data Dictionary Codebook

12-10-2023 1:37am

| Languages | | |
|---|---|---|
| ID | | Display Name |
| en | ☑ | English **(default)** |
| ur | ☐ | urdu |

| # | Variable / Field Name | Field Label<br>*Field Note* | Field Attributes (Field Type, Validation, Choices, Calculations, etc.) |
|---|---|---|---|
| **Instrument: Questionnaire (questionnaire)** Enabled as survey<br>Active languages - Data Entry: en, ur \| Survey: en, ur | | | |
| 1 | [participant_id] | Participant ID | text |
| 2 | [questionnaire] | I have read and understand the above consent form. I certify that I am 18 years old or older. By clicking the "Next Page" button to enter the survey, I indicate my willingness to voluntarily take part in this study. | descriptive |
| 3 | [district] | Section Header: *Demographics*<br>District | text |
| 4 | [age] | Age | radio, Required<br>1 18-25<br>2 26-34<br>3 35-44<br>4 45-54<br>5 55-64<br>6 65+ |
| 5 | [gender] | Gender | radio, Required<br>1 Male<br>2 Female<br>3 I prefer not to say |
| 6 | [designation] | What is your Designation/ official position? | radio, Required<br>1 Veterinary Technician<br>2 AI Technician<br>3 No Certification |
| 7 | [practice_type] | Practice type | radio, Required |

| | | | |
|---|---|---|---|
| | | | 1 Field practice<br>2 Clinic<br>3 Both |
| 8 | [experience] | For how many years have you been practicing in this position? | radio, Required<br>1 1-5<br>2 6-10<br>3 11-20<br>4 21-25<br>5 26-30<br>6 30+ |
| 9 | [cases_handeld] | How many cases of vector-borne diseases do you see per month? | text, Required |
| 10 | [attended_seminar_tbd] | Section Header: *Knowledge*<br>Have you ever attended seminars/workshops/training about ticks and tick-borne diseases? | yesno, Required<br>1 Yes<br>0 No |
| 11 | [frequency_of_attending_seminar] | What frequency have you attended seminars/workshops/training about ticks and tick-borne diseases? | radio, Required<br>1 Twice a year<br>2 Once a year<br>3 Once in your career<br>4 Never |
| 12 | [common_tick] | What is the most common tick species in your area? | text |
| 13 | [tick_on_animal_body] | Where on the animal's body do ticks commonly bite and attach? | radio, Required<br>1 Neck<br>2 Chest<br>3 Inner Thighs<br>4 Perineum<br>5 Udder<br>6 External of genitalia<br>7 Ears<br>8 Dewlap<br>9 other |
| 14 | [animal_with_more_tbd] | From your experience, which animal species is more affected by ticks? | radio, Required<br>1 Cattle<br>2 Buffalo<br>3 Sheep |

| | | | |
|---|---|---|---|
| | | | 4 Goat<br>5 Dogs and Cats<br>6 Others |
| 15 | [habitat_of_tick] | What is the preferred habitat of ticks? | radio, Required<br>1 Cracks or Crevices<br>2 Forest Areas<br>3 Grassland<br>4 Arid deserts |
| 16 | [tbd_zoonotic] | Can ticks transmit disease from animal to human? | yesno, Required<br>1 Yes<br>0 No |
| 17 | [name_zoonotic_tbd] | Can you name any disease transmitted from animals to humans through ticks? | text |
| 18 | [tick_attachment_duration] | How long does a tick have to be attached to the skin to transmit a disease? | radio, Required<br>1 Days<br>2 Hours<br>3 Minutes<br>4 Seconds |
| 19 | [tick_bite_identify] | Does a person always know when a tick has bitten them? | yesno, Required<br>1 Yes<br>0 No |
| 20 | [heard_of_any_of_tbd] | Have you heard of any of these tick-borne diseases of animals? (Select Multiple) | radio, Required<br>1 Babesiosis<br>2 Theileriosis<br>3 Anaplasmosis<br>4 Ehrlichiosis<br>5 Rickettsiosis<br>6 Q-fever<br>7 None of these |
| 21 | [endemic_tbd] | Which of these tick-borne diseases of animals is endemic to your area? | radio, Required<br>1 Babesiosis<br>2 Theileriosis<br>3 Anaplasmosis<br>4 Ehrlichiosis<br>5 Rickettsiosis<br>6 Q-fever |

| | | | |
|---|---|---|---|
| | | | 7 None of these |
| 22 | [name_tbd_human] | Can you name any tick-borne diseases of humans in your area? | text |
| 23 | [how_will_you_rank_ticks_an] | How will you rank ticks and tick-borne diseases as animal health problems in your area? | slider (number, Min: 0, Max: 100), Required<br>Slider labels: Not a problem, Mild problem, Big problem<br>Custom alignment: RH |
| 24 | [would_you_like_to_receive] | Section Header: *Attitude*<br>Would you like to receive training on ticks and tick-borne diseases? | radio, Required<br>1 Yes<br>2 No<br>3 Unsure |
| 25 | [format_of_training] | What format would you prefer for the training you want to receive? | radio, Required<br>1 In-person seminars<br>2 Online modules<br>3 Written instruction<br>4 Other |
| 26 | [farmers_act_on_your_advice] | Do you think farmers act on your advised strategies of tick management and control in their herd? | yesno, Required<br>1 Yes<br>0 No |
| 27 | [farmers_benfited_by_tick_control] | Do you feel farmers would benefit from more tick control practices? | yesno, Required<br>1 Yes<br>0 No |
| 28 | [barriers_in_tick_control] | Which of the following do you think are barriers to farmers following tick control? | radio, Required<br>1 Cost<br>2 Time<br>3 Access to acaricides/chemicals<br>4 Lack of concern |
| 29 | [farmers_at_risk] | Do you feel farmers are at risk of a tick bite? due to their practices | yesno, Required<br>1 Yes<br>0 No |
| 30 | [tbd_threat_animal_your_area] | Do you think the tick-borne disease is a concern for animals in your area | yesno, Required<br>1 Yes<br>0 No |
| 31 | [tbd_threat_to_human_your_area] | Do you think the tick-borne disease is a concern for people in your area | yesno, Required<br>1 Yes<br>0 No |

| | | | |
|---|---|---|---|
| 32 | [use_diagnostics_test] | Section Header: *Practices*<br>Do you use diagnostic tests for any tick-borne diseases? | yesno, Required<br>1 Yes<br>0 No |
| 33 | [which_diagnostic_tests] | Which diagnostic tests for any tick-borne diseases? | text |
| 34 | [treated_for_tbd_by_u] | Have you ever treated animals for any tick-borne disease? | yesno, Required<br>1 Yes<br>0 No |
| 35 | [which_treatment_practices] | Which treatment practices do you use for the treatment of tick-borne diseases? | text |
| 36 | [educate_farmers_tbd] | Do you educate farmers about ticks and tick-borne diseases? | yesno, Required<br>1 Yes<br>0 No |
| 37 | [losses_farmer_tbd_peryear] | How many losses does a farmer face per year due to tick-borne disease? (PKR) | text |
| 38 | [tick_control_recommendation] | Which kind of tick control and management practices do you recommend to farmers for their animals? (Select all that apply) | radio, Required<br>1 Acaricide use<br>2 Deworming<br>3 Biosecurity measures<br>4 Herbal<br>5 None |
| 39 | [best_practice_personal_protection] | What is the best practice to protect ourselves against ticks? | text |
| 40 | [check_yourself_after_livestock] | Do you check yourself for ticks after visiting the livestock farms? | yesno, Required<br>1 Yes<br>0 No |
| 41 | [ever_bitten_by_tick] | Have you ever been bitten by a tick? | yesno, Required<br>1 Yes<br>0 No |
| 42 | [fever_after_bite] | Have you ever observed fever-like symptoms after tick-bite? | yesno, Required<br>1 Yes<br>0 No |
| 43 | [farmer_consult_after_bite] | Do farmers consult with you after ticks bite them or their farm-worker s? | yesno, Required<br>1 Yes<br>0 No |
| 44 | [questionnaire_complete] | Section Header: *Form Status*<br>Complete? | dropdown<br>0 Incomplete |

| | | | |
|---|---|---|---|
| | | | 1 Unverified<br>2 Complete |
