## Supplementary material for "Knowledge, attitudes, and practices of para-veterinary workers about ticks and tick-borne diseases in three provinces of Pakistan": Scoring criteria

### Demographics

| Questions | Responses | Total Score |
|---|---|---|
| Province | | |
| District | | |
| Age | 18-25 | |
| | 26-34 | |
| | 35-44 | |
| | 45-54 | |
| | 55-64 | |
| | 65+ | |
| Gender | Female | |
| | Male | |
| | I prefer not to say | |
| What is your Designation/ official position? | Veterinary Technician | |
| | AI Technician | |
| | No Certification | |
| Practice type | Field practice | |
| | Clinic | |
| | Both | |
| For how many years have you been practicing in this position? | 1-5 | |
| | 6-10 | |
| | 11-20 | |
| | 21-25 | |
| | 26-30 | |
| | 30+ | |
| How many cases of vector-borne diseases do you see per month? | | |

### Workshop Attendance

| Questions | Responses | Total Score |
|---|---|---|
| Have you ever attended seminars/workshops/training about ticks and tick-borne diseases? | Yes | |
| | No | |
| What frequency have you attended seminars/workshops/training about ticks and tick-borne diseases? | Twice a year | |
| | Once a year | |
| | Once in your career | |
| | Never | |

### Knowledge

| Questions | Responses | Total Score |
|---|---|---|
| What is the most common tick species in your area? | NA/ Other Tick =0<br>Hyalomma =1<br>Rhipicephalus =1<br>Hyalomma + Rhipicephalus =2 | 2 |
| Where on the animal's body do ticks commonly bite and attach? | Neck=1 | 1 |
| | Chest=1 | |
| | Inner Thighs=1 | |
| | Perineum=1 | |
| | Udder=1 | |
| | External of genitalia=1 | |
| | Ears=1 | |
| | Dewlap=1 | |
| | Don't Know=0 | |
| From your experience, which animal species are more affected by ticks? | Cattle=2 | 2 |
| | Buffalo=2 | |
| | Sheep=1 | |
| | Goat=1 | |
| | Dogs and Cats=1 | |
| | Others=0 | |
| What is the preferred habitat of ticks? | Cracks or Crevices=1 | 1 |
| | Forest Areas=1 | |
| | Grassland=1 | |
| | Arid deserts=0 | |
| Can ticks transmit disease from animal to human? | Yes=1 | 1 |
| | No=0 | |
| Can you name any disease transmitted from animals to humans through ticks? | Correct disease name =1<br>NA/Wrong name =0 | 1 |
| How long does a tick have to be attached to the skin to transmit a disease? | Days = 1 | 1 |
| | Hours = 0 | |
| | Minutes = 0 | |
| | Seconds = 0 | |
| Does a person always know when a tick has bitten them? | Yes =0 | 1 |
| | No =1 | |
| Have you heard of any of these tick-borne diseases of animals? | Babesiosis=1 | 1 |
| | Theileriosis=1 | |
| | Anaplasmosis=1 | |
| | Ehrlichiosis=1 | |
| | Rickettsiosis=1 | |
| | Q-fever=1 | |
| | No=0 | |
| Which of these tick-borne diseases of animals is endemic to your area? | Babesiosis=1 | 1 |
| | Theileriosis=1 | |
| | Anaplasmosis=1 | |
| | Ehrlichiosis=1 | |

| | | |
|---|---|---|
| | Rickettsiosis=1<br>Q-fever=1<br>None=0<br>I Don't Know=0 | |
| Can you name any tick-borne diseases of humans in your area? | Correct = 1<br>NA/wrong =0 | 1 |
| How will you rank ticks and tick-borne diseases as animal health problems in your area? | 1. Not a problem<br>2. Mild problem<br>3. Big problem | |

**Attitude**

| Questions | Responses | Total Score |
|---|---|---|
| Would you like to receive training on ticks and tick-borne diseases? | Yes =1<br>No = 0<br>Unsure = 0 | 1 |
| What format would you prefer for the training you want to receive? | In-person seminars=1<br>Online modules=1<br>Written instruction=1<br>Other=0 | 1 |
| Do you think farmers act on your advised strategies of tick management and control in their herd? | Yes=1<br>No=0 | 1 |
| Do you feel farmers would benefit from more tick control practices? | Yes = 1<br>No = 0 | 1 |
| Do you feel farmers are at risk of a tick bite? due to their practices | Yes=1<br>No=0 | 1 |
| Do you think the tick-borne disease is a concern for animals in your area | Yes = 1<br>No = 0 | 1 |
| Do you think the tick-borne disease is a concern for people in your area | Yes=1<br>No=0 | 1 |
| Which of the following do you think are barriers to farmers following tick control? | Cost<br>Time<br>Access to acaricides/chemicals<br>Lack of concern | |

**Practices**

| Questions | Responses | Total Score |
|---|---|---|
| Do you use diagnostic tests for any tick-borne diseases? | Yes=1<br>No=0 | 1 |

| | | |
|---|---|---|
| Which diagnostic tests for any tick-borne diseases? | Blood test =1<br>All other = 0 | 1 |
| Have you ever treated animals for any tick-borne disease? | Yes=1 | 1 |
| | No=0 | |
| Which treatment protocol do you use for the treatment of tick-borne diseases? | Anti-tick-borne disease medicine =1<br><br>NA/ other =0 | 1 |
| Do you educate farmers about ticks and tick-borne diseases? | Yes =1 | 1 |
| | No = 0 | |
| Which kind of tick control and management practices do you recommend to farmers for their animals? | Acaricide use = 1 | 1 |
| | Deworming = 1 | |
| | Biosecurity measures = 1 | |
| | Herbal = 0 | |
| | None = 0 | |
| What is the best practice to protect ourselves against ticks? | Any personal protective measure=1<br>NA/Other =0 | 1 |
| Do you check yourself for ticks after visiting the livestock farms? | Yes=1 | 1 |
| | No=0 | |
| Do farmers consult with you after ticks bite them or their farm workers? | Yes=1 | 1 |
| | No=0 | |
| How many losses does a farmer face per year due to tick-borne disease? (PKR) | | |
| Have you ever been bitten by a tick? | Yes | |
| | No | |
| Have you ever observed fever-like symptoms after tick-bite? | Yes | |
| | No | |

| **Category** | **Total Score** |
|---|---|
| **Knowledge** | **13** |
| **Attitudes** | **7** |
| **Practices** | **9** |
